# Elucidating the diversity and potential function of nonribosomal peptide and polyketide biosynthetic gene clusters in the root microbiome

**DOI:** 10.1101/2020.06.07.138487

**Authors:** Barak Dror, Zongqiang Wang, Sean F. Brady, Edouard Jurkevitch, Eddie Cytryn

**Author notes:** Address correspondence to Eddie Cytryn.

## Abstract

Polyketides (PKs) and nonribosomal peptides (NRPs) are two microbial secondary metabolite (SM) families known for their variety of functions, including antimicrobials, siderophores and others. Despite their involvement in bacteria-bacteria and bacteria-plant interactions, root-associated SMs are largely unexplored due to the limited cultivability of bacteria. Here, we analyzed the diversity and expression of SM-encoding biosynthetic gene clusters (BGCs) in root microbiomes by culture-independent amplicon sequencing, shotgun metagenomics and metatranscriptomics. Roots (tomato and lettuce) harbored distinct compositions of nonribosomal peptide synthetases (NRPSs) and polyketide synthases (PKSs) relative to the adjacent bulk soil, and specific BGC markers were both enriched and highly expressed in the root microbiomes. While several of the highly abundant and expressed sequences were remotely associated with known BGCs, the low similarity to characterized genes suggests their potential novelty. Low similarity genes were screened against a large set of soil-derived cosmid libraries, from which five whole BGCs of unknown function were retrieved. Three clusters were taxonomically affiliated with Actinobacteria, while the remaining were not associated with known bacteria. One Streptomyces-derived BGC was predicted to encode for a polyene with potential antifungal activity, while the others were too novel to predict chemical structure. Screening against a suite of metagenomic datasets revealed a higher abundance of retrieved clusters in roots and soil samples. In contrast, they were almost completely absent in aquatic and gut environments, supporting the notion that they might play an important role in root ecosystems. Overall, our results indicate that root microbiomes harbor a specific assemblage of undiscovered SMs.

**Importance:** We identified distinct secondary metabolite (polyketide and nonribosomal peptide) encoding genes that are enriched (relative to adjacent bulk soil) and expressed in root ecosystems, yet almost completely absent in human gut and aquatic environments. Several of the genes were distantly related to genes encoding for antimicrobials and siderophores, and their high sequence variability relative to known sequences suggests that they may encode for novel metabolites and may have unique ecological functions. This study demonstrates that plant roots harbor a diverse array of unique secondary metabolite encoding genes that are highly enriched and expressed in the root ecosystem. The secondary metabolites encoded by these genes might assist the bacteria that produce them in colonization and persistence in the root environment. To explore this hypothesis, future investigations should assess their potential role in inter-bacterial and bacterial-plant interactions.

## Introduction

Soil is an extremely diverse ecosystem that contains a myriad of micro- and macro-organisms, including nematodes, arthropods, fungi and bacteria. The rhizosphere is a narrow region of soil directly influenced by root exudates and mucilage (1, 2). This “hot spot” of organic matter and nutrients “enriches” a specific fraction of the soil microbial community known as the root microbiome, which is significantly different than the surrounding soil microbiome (3). Over the past two decades, several studies have linked specific constituents of the root microbiome to enhanced plant growth and development and inhibition of soilborne plant pathogens (4), either by direct antagonism, and/or induced systemic resistance (5). These functions are often facilitated by the vast array of secondary metabolites (SMs) produced by root-associated bacteria, which play a key role in inter- and intra-species interactions (6, 7).

Many important soil and root-associated bacterial SMs are nonribosomal peptides (NRPs), or polyketides (PKs), produced by nonribosomal peptide synthetases (NRPSs) or polyketide synthases (PKSs), respectively. These are encoded on large biosynthetic gene clusters (BGCs) that often exceed 50,000 bp (8). Enzymatic complexes in these families follow a similar biosynthetic logic wherein molecules are assembled in an iterative building process using conserved domains that are organized in modules (9, 10). NRPSs and PKSs are responsible for the synthesis of a wide array of siderophores, toxins, pigments and antimicrobial compounds (11) that are believed to play a pivotal role in bacterial adaptation to soil and rhizosphere ecosystems, and in plant health and development (12). Despite their ecological (rhizosphere competence) and translational (biocontrol agents and novel antimicrobial compounds for plant protection) importance, little is known about the occurrence, diversity and dynamics of NRPSs and PKSs in root ecosystems, or their role in intra- and inter-microbial and plant-bacterial interactions.

A major challenge in exploring the role and function of SMs in soil stems from the fact that the majority of bacteria cannot be cultivated using conventional methods, making it difficult to study these bacteria and the diversity, expression and function of the metabolites they produce (13). Despite progress made in culturing techniques, our capacity to isolate soil and root-associated bacteria is highly constrained, primarily because it is challenging to mimic the natural conditions required for growing these bacteria (14). Furthermore, many bacterial BGCs are silent under laboratory conditions and therefore the metabolites that they encode are extremely challenging to isolate (15).

To circumvent the above mentioned barriers, a myriad of culture-independent sequencing-based and omics tools have been developed to reveal the scope and composition of soil-derived BGCs encoding NRPSs and PKSs (16, 17), and to infer the chemical composition and structure of the metabolites produced by these synthases (18, 19). For instance, amplicon sequencing-based approaches have been developed to target short fragments within adenylation (AD-in NRPS) and ketosynthase (KS-in PKS) domains. These amplicons can be used to ascertain the diversity and abundance of bacterial BGCs in complex environments, as both AD and KS domains are important (in concert with other components) for the assembly, and thus identity and activity, of the synthesized metabolites (20, 21). To date, a handful of studies have explored the diversity and composition of bacterial SM-encoding BGCs in soil, demonstrating the vast genetic diversity and novelty of NRPSs and PKSs genes (22, 23). However, little is known regarding the distribution and of these gene families in the root microbiome, and their functional role in this complex community remains an enigma (24).

This study proposes a unique approach to analyze the diversity and potential functions of NRPSs and PKSs in the root, specifically focused on elucidating: (i) the composition and diversity of NRPSs and PKSs encoding genes in the root environment relative to adjacent bulk soil; (ii) NRPSs and PKSs composition and expression in the root as a function of plant type; (iii) the sequence and inferred SM structures of whole bacterial BGCs that are highly abundant or expressed in root environments; and (iv) the occurrence of root-enriched bacterial BGCs in other ecosystems.

## Results

### 1. Composition and diversity of NRPSs and PKSs genes in roots vs. bulk soil

To determine the composition and diversity of NRPSs and PKSs genes in tomato and lettuce root samples relative to bulk soil (previous studies targeting this controlled lysimeter system showed that bulk soil from tomato and lettuce microbiomes were almost identical and therefore only tomato soil was analyzed here), we applied a previously described amplicon sequencing approach to amplify the conserved adenylation (AD) and ketosynthase (KS) domains of NRPSs and PKSs, respectively (23). Overall, sequencing yielded a total of 1,850,442 and 2,174,020 raw KS and AD reads with average read lengths of 280 bp and 235 bp, respectively (**Table S1**). Further filtering steps using QIIME2 and DADA2 denoising methods, resulted in 2,980 and 3,269 non-redundant KS and AD domain sequences, respectively.

We observed significantly higher diversity of both AD and KS domains in the bulk tomato soil compared to the adjacent roots (**Fig. S1-A and C**). In contrast, no difference in diversity was observed between tomato and lettuce roots for either of the SM encoding domains (**Fig. S1-B and D**). To assess differences in AD and KS domain diversity between samples, a principal coordinate analysis (PCoA) and analysis of similarity (ANOSIM) using the Bray-Curtis similarity index were performed (Fig. **1A** and **1B**). The KS and AD domain profiles of the roots (from both tomato and lettuce) formed distinct clusters, which were significantly different from the adjacent bulk soil (R=0.332, p< 0.05 for PKS, R=0.308, p<0.01 for NRPS).

**Figure 1.**
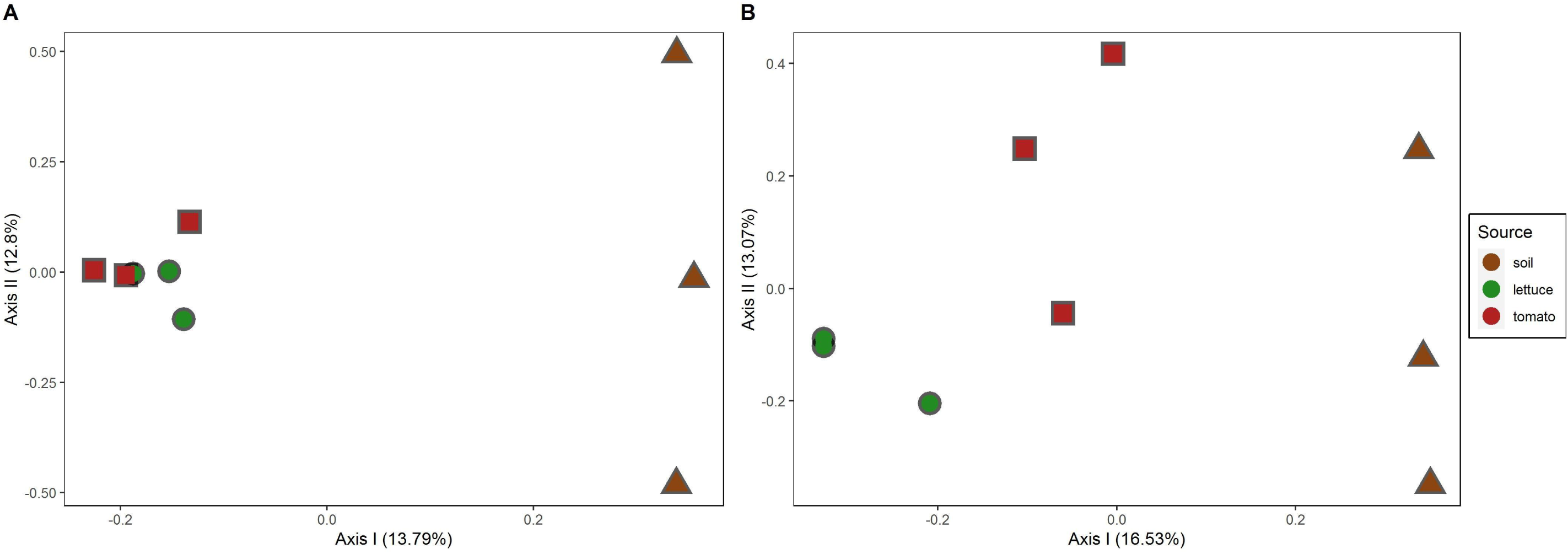
Principal coordinate analysis (PCoA) of lettuce roots (green circles), tomato roots (red squares) and soil (from tomato lysimeters, brown triangles) samples using the Bray-Curtis similarity index. (A) Analysis of KS (from PKS) domain amplicons (roots vs. soil ANOSIM statistic R=0.332, p-Value<0.05, tomato vs. lettuce vs. soil ANOSIM R=0.432, NS). (B) Analysis of AD (NRPS) domain amplicons (Roots vs. soil ANOSIM statistic R=0.308, p-Value<0.01, tomato vs. lettuce vs. soil ANOSIM R=0.6872, p-Value<0.01).

To explore the potential novelty of root-associated SM-encoding genes, the amplified AD and KS domain sequences from the root and soil samples were first aligned against the MIBiG (Minimum Information about a Biosynthetic Gene cluster) repository (25) using blastp, with >50% amino acid sequence identity cut-off, and then grouped according to their identity to the MIBiG reference genes (**Table 1**). On average, more than 25% of the AD and almost 13% of the KS domain sequences in the root environment showed less than 50% amino acid identity with genes found in the MIBiG database (characterized as ‘unassigned’), whereas less than 1% and 6% of the AD and KS sequences (respectively) shared over 85% similarity to the reference MIBiG genes. These results demonstrate the profusion of potentially novel SMs in both root and soil environments.

**Table 1.** Abundance of NRPS (AD) and PKS (KS) amplicons based on levels of similarity to MIBiG NRPSs and PKSs reference genes, in the tomato and cucumber root and tomato soil samples. Identity groups were determined as unassigned (< 50% amino acid sequence identity), 50-70%, 70-85%, 85-95% and 95-100% sequence identity. Numbers outside parentheses indicate % of sequences associated with each group, numbers inside parentheses indicate total number of hits for each group.

### 2. Pinpointing predicted root-enriched NRPs and PKs

As SMs are known to play critical roles in bacteria-bacteria and bacteria-plant interactions, we were interested in the associated metabolites synthesized by BGCs whose AD or KS domains were highly abundant and enriched in tomato or lettuce roots relative to bulk tomato soil. To do so, the MIBiG-aligned amplicons were annotated to corresponding BGC-associated metabolites, using a cut-off e-Value of < 10^−40^. Sequences that did not meet these criteria were defined as ‘unknown’. Previous analyses have shown that when compared to reference KS or AD domain sequences, amplicons with e-Values of < 10^−40^ are likely derived from the same BGC family as the reference sequence, and thus may be inferred to encode a similar function (26-29).

A differential abundance analysis using DESeq2 of the top20 highly abundant AD and KS amplicons revealed that 55% (11/20) and 70% (14/20) of the amplicons in both of the plant root samples (tomato and lettuce respectively, adjusted p-value < 0.1, log2FoldChange>5), were not associated with known BGCs, thus their associated metabolites cannot be inferred (**Fig. 2**). Nonetheless, several root-enriched AD and KS domains sequences (9 in tomato and 6 in lettuce) were above these threshold values, thus can be considered as congener to known metabolites. These included the non-ribosomal peptides stenothricin (30) and griselimycin (31) whose BGC analogues were highly abundant in the tomato and lettuce roots, respectively, and were less profuse in the bulk soil. While we cannot determine the actual role of these enriched metabolites in the root environment, both of them are known for their antimicrobial activity.

**Figure 2.**
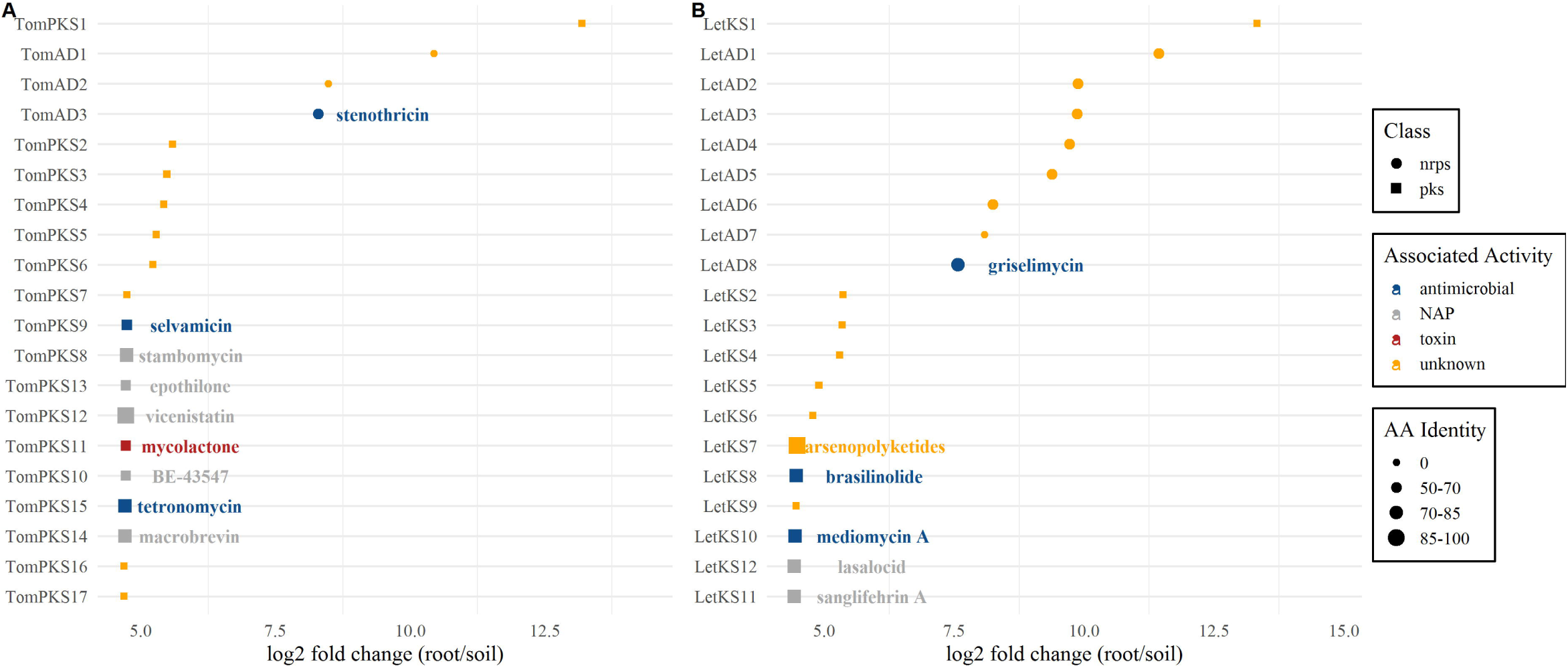
Differential abundance of the top20 highly abundant NRPS and PKS amplicons in tomato (A) and lettuce (B) root microbial communities relative to bulk tomato soil. DESeq 2 was used to calculate log2 fold changes and test for significance (adjusted p-value (FDR) of <0.1). Associated metabolites (based on the MIBiG repository) are presented only for sequences with e-Value<10-40 and >50% AA identity. Identity bars represent amino acid identity to MIBiG reference sequence (0-not identified).

Next, we calculated the relative abundance of amplicons that were associated with known metabolites (based on the criteria above) in the different root and soil samples (**Fig. S2**).To pinpoint associated metabolites that may play a role in adaptation to the root environment, we focused our analysis on metabolites that were present in at least four of the root samples (tomato and lettuce) and in no more than one soil sample (**Fig. 3**). In addition, to identify metabolites specifically relevant to soil, we also selected metabolites that were present in all three soil samples and in no more than one root-associated sample. For NRPs, we found four associated metabolites that were highly abundant in both of the root samples (*e*.*g*. the Streptomyces-derived antibiotic macrolide family streptovaricin). For PKs, we again found several highly abundant Streptomyces-derived predicted metabolites, among others. These included lasalocid, sanglifehrin A and azalomycin A. Interestingly, amplicons associated with the two former were also found to be highly enriched in roots relative to soil in lettuce (Fig. 2B). Overall, we found 26 associated metabolites were present in at least one of the root-associated samples, and completely absent in the soil samples, *e*.*g*. diaphorin (in lettuce) and basiliskamides (present in both tomato and lettuce, **Fig. S2**).

**Figure 3.**
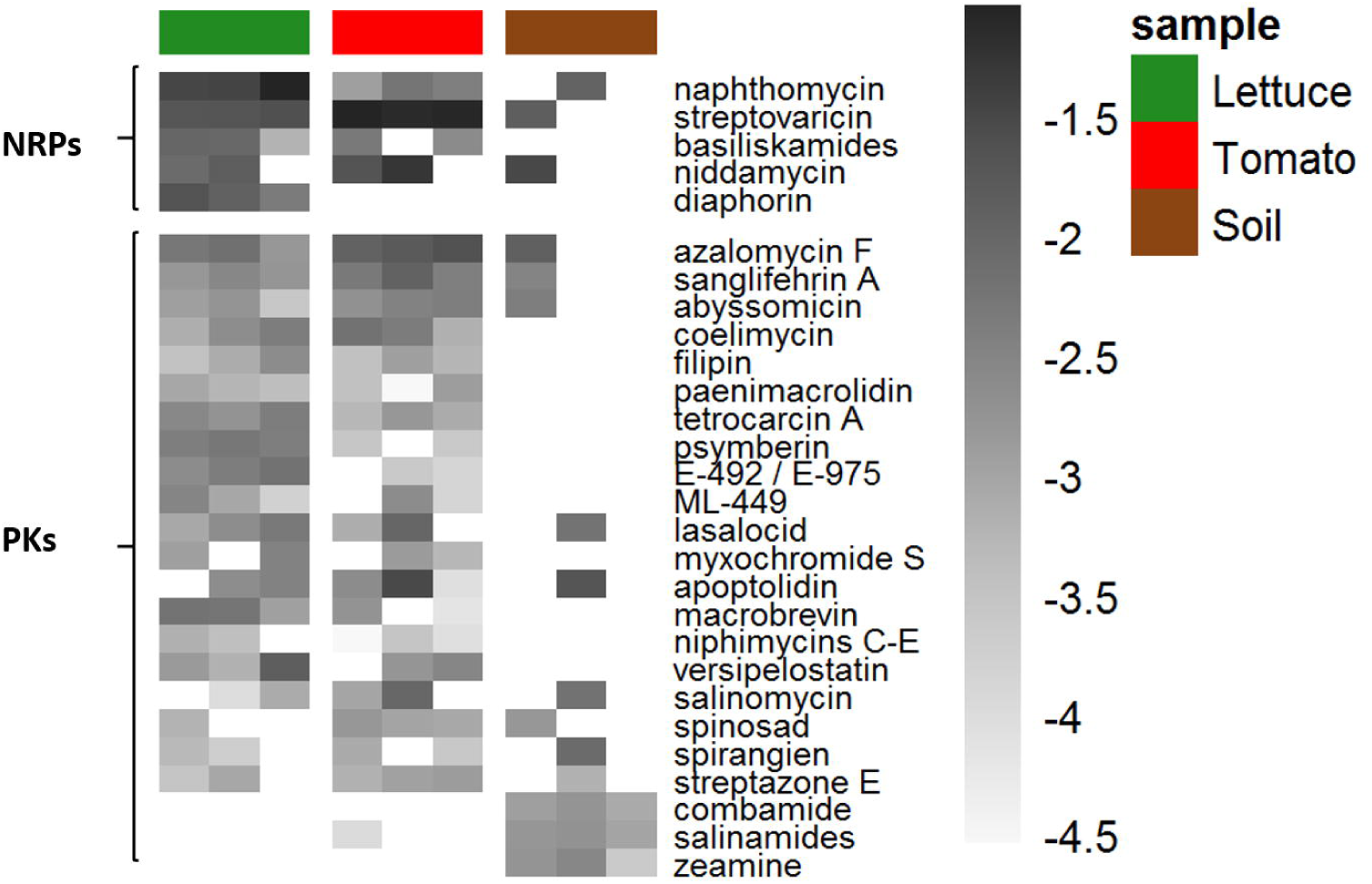
Relative abundance of NRPS (A) and PKS (B) associated metabolites based on MIBiG annotations. Only AD and KS amplicon hits to reference MIBiG sequences over >50% amino acid identity and e-Value<10-40 were included. Counts were normalized and log10-transformed. Only associated metabolites that were present in >2 different samples are shown.

Due to the potential biases associated with the above PCR-based approach, we analyzed previously published (32) shotgun metagenomes of the same tomato and lettuce root samples (n=3 each). Assembled open reading frame (ORF) sequences from the metagenomes identified using Prodigal were aligned against the MIBiG database, generating a list of the fifty most abundant NRPSs and PKSs in each of the root datasets (representing the normalized abundance within-samples by plotting the coefficient of variance (CV) for each gene, **Fig. S3A** and **B**). In addition, as gene clusters are often silent or expressed under very specific conditions, we evaluated gene expression in parallel to gene occurrence to uncover active BGCs with ecological importance in the highly dynamic root ecosystem. Thus, in parallel to the shotgun metagenome analysis described above we applied a similar analytical approach using the previously collected shotgun meta-transcriptomics (32) to identify NRPSs and PKSs with enhanced expression in lettuce and/or tomato root microbiomes. Interestingly, 60% (30/50) and 46% (23/50) of the AD and KS domains that were highly abundant in the tomato and lettuce root samples (respectively) were highly expressed as well (**Fig. S**3).

Next, we filtered the highly abundant hits based on their CV value (<50) in order to analyze sequences with lower dispersion levels within tomato and lettuce root-associated samples, followed by taxonomical annotation using MEGAN. The resulting 42 sequences were clustered into two main phyla-Actinobacteria (13/42) and Proteobacteria (25/42). Several sequences could not be assigned taxonomic affiliation and one was assigned to the Bacteriodetes phylum (**Fig. 4**). While we could not infer the associated metabolites of most of these highly abundant sequences (including all of those assigned to the phylu Proteobacteria), suggesting their potential novelty, we did manage to annotate several of the Actinobacteria-associated metabolites. These were distantly associated with Ossamycin (5 hits) and Polycyclic Tetramate Macrolactams (PTMs; 2 hits, including the most highly abundant NRPSs/PKSs-related sequence and the 5^th^ most expressed). Ossamycin is a known antifungal and cytotoxic macrocyclic polyketide originally isolated from soil-associated Streptomyces hygroscopicus var. ossamyceticus (33, 34), while PTMs are a family of biologically important metabolites including HSAF, ikarugymcin and clifednamides, generally associated with different isolates of Actinobacteria and Gammaproteobacteria (35).

**Figure 4.**
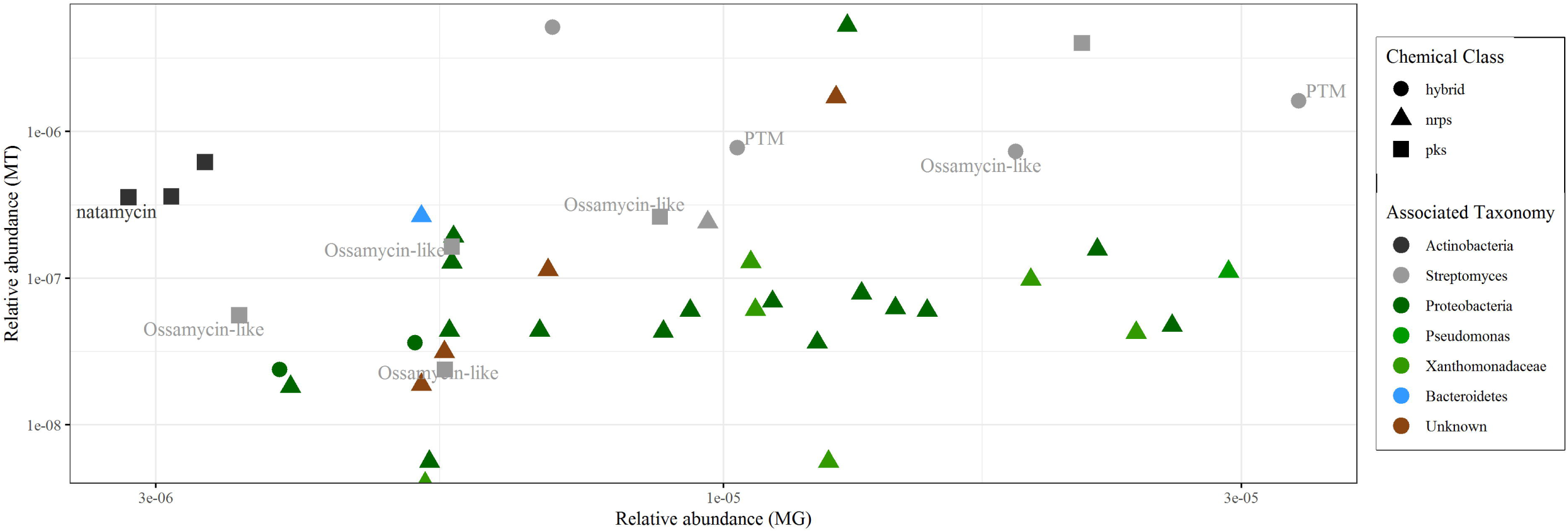
Relative abundance and taxonomy of highly abundant NRPSs/PKSs in tomato and lettuce roots metagenome (MG) and metatranscriptome (MT). Shapes represent BGC-associated chemical class. Taxonomy was assigned using MEGAN 5.10 and the lowest common ancestor (LCA) classification algorithm. Only low CV sequences were chosen.

Finally, to evaluate the extent to which the amplicon sequencing method was able to detect NRPSs and PKSs in the targeted samples relative to the PCR-independent shotgun-metagenomic analyses, we analyzed the distribution of total MIBiG-associated genes in all four datasets (lettuce and tomato NRPSs and PKSs amplicons, tomato and lettuce metagenomes, **Fig. S4**). In general, approximately 34% and 25% of the MIBiG-characterized genes were found in both amplicon sequences and metagenomes of the lettuce and tomato roots, respectively. In contrast, approximately 33% and 20% of the tomato and lettuce genes, respectively, were only detected in the shotgun metagenomic datasets. Less than 4% and 5% of the NRPSs and PKSs were found only within the tomato and lettuce amplicon sequencing datasets (28 and 35 genes in tomato and lettuce, respectively) and less than a fifth (139 genes or 17.2%) were common to all four culture-independent datasets.

## 3. Extraction and environmental distribution of whole SM-encoding gene clusters

The identification of NRPSs and PKSs that were either enriched in roots relative to adjacent bulk soil or were abundant and/or highly expressed in lettuce and/or tomato roots, encouraged us to capture whole BGCs associated with these sequences, in order to shed light on their phylogenetic affiliation and potentially infer their function and chemical structure. This was achieved by screening the root-associated NRPSs and PKSs candidate sequences identified in this study against a large set of previously collated soil and rhizosphere cosmid libraries using the bioinformatic platform eSNaPD (29)(see Materials and Methods for full pipeline). Five cosmid library targets showed low e-Values and high nucleotide sequence identity (>75%) to candidate AD or KS sequences. Sequencing and annotation of the metagenomic insert captured in each cosmid revealed three NRPS and two hybrid NRPS/PKS gene clusters (**Fig. 5**). Based on gene content and sequence identity, the identified gene clusters were not identical to BGCs associated with known metabolites. The NRPS and PKS ORFs of two recovered clones (B326 and B385) were not affiliated with any known bacterial taxa (<50% nucleic acid identity to the NCBI database), while the other three clones were related to genes from Actinobacteria **(Table 2)**.Of the cosmids recovered from the metagenomic libraries, clone B481 was nearly identical to an uncharacterized NRPS BGC found in the genome of *Streptomyces cyaneogriseus* (**Fig. 6A**). The only predicted chemical structure we could infer from the recovered BGCs was for clone B893, which was related to an uncharacterized PKS gene cluster found in the genome of *Saccharothrix saharansis*, a filamentous Actinobacteria isolated from desert soil. A detailed bioinformatic analysis of its PKS domains revealed that the gene cluster likely encodes for an extended polyene substructure (**Fig. 6B**). The seven PKS modules captured on the clone all contain dehydratase (DH) and ketoreduction (KR) domains indicating that each module introduces a double bond into the polyketide backbone (**Fig. 6B**). While polyene substructures like this are seen in a number of natural products (36, 37), they are most commonly seen in polyene antifungal agents, including many that are derived from *Streptomyces* species (e.g., cyphomycin, nystatin, filipin and pimaricin). This may suggest that the BGC encode for an antifungal compound.

**Figure 5.**
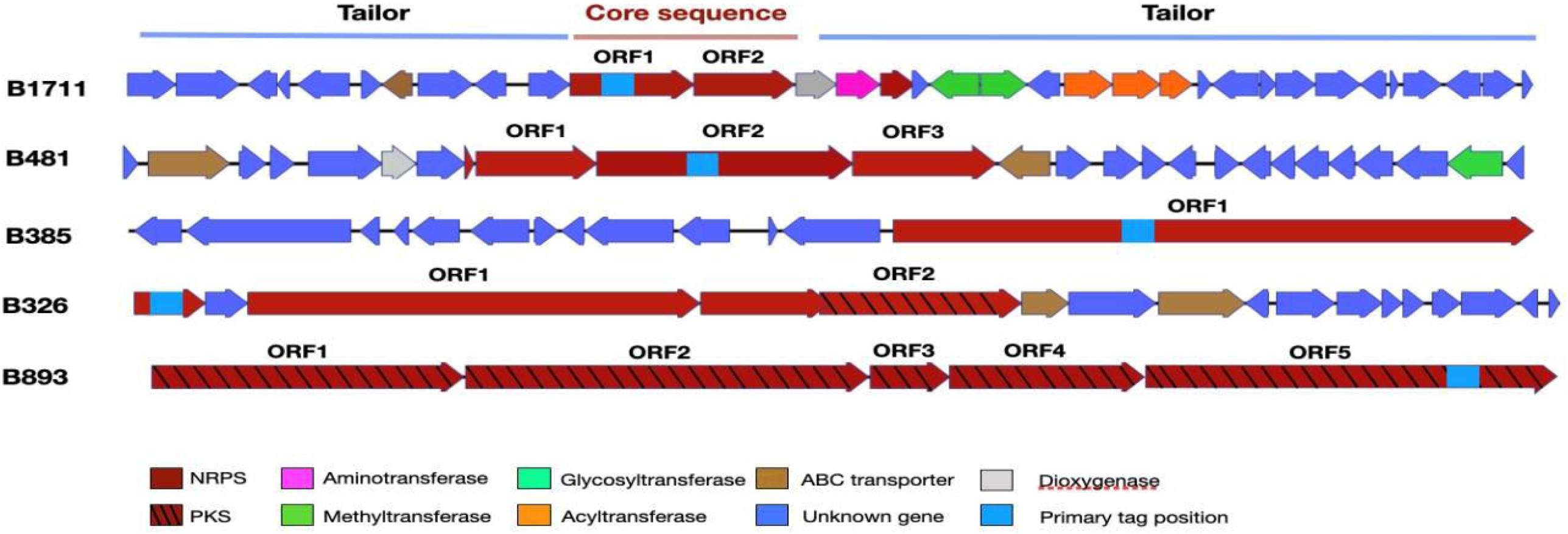
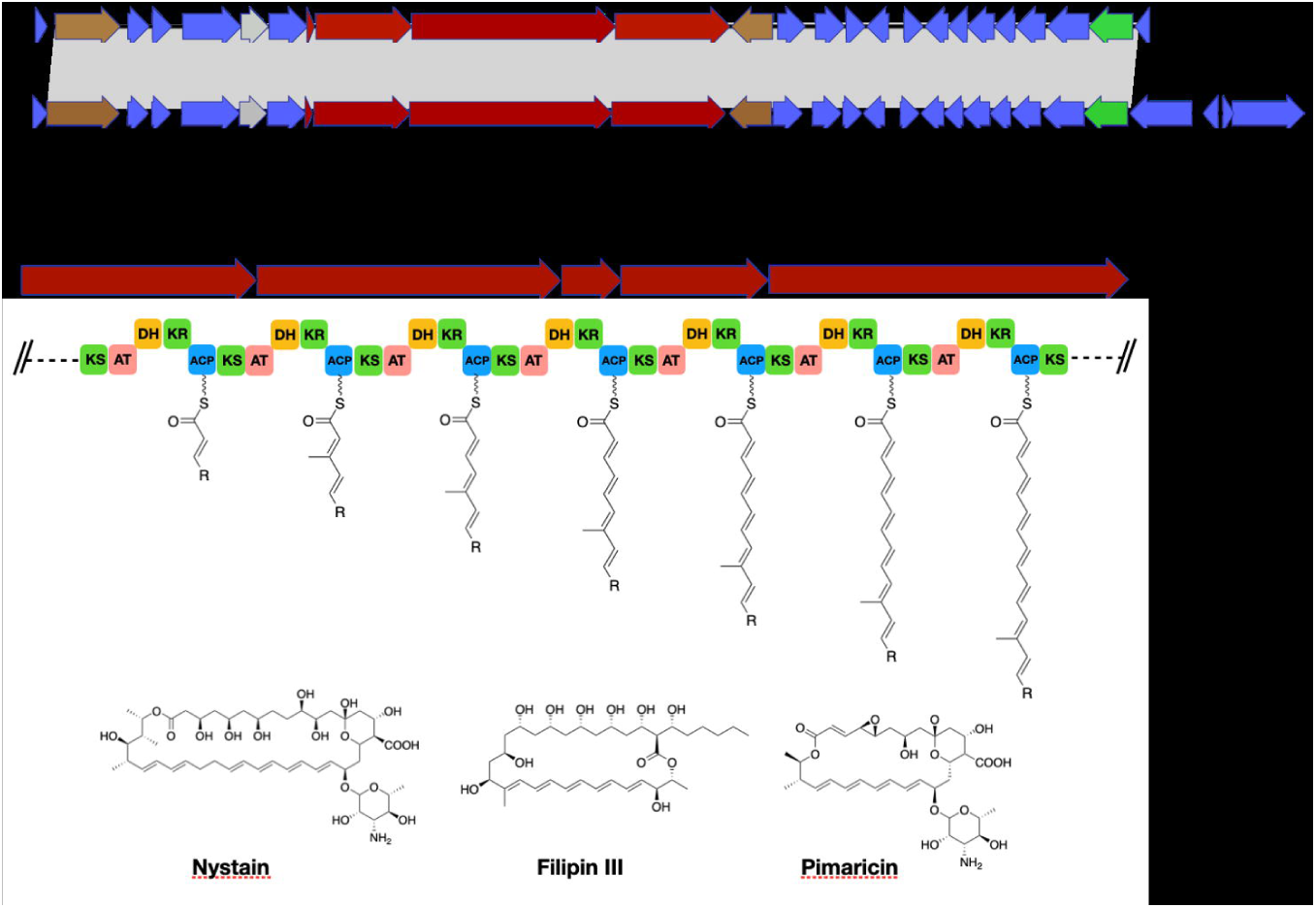
Overview of retrieved (from archived soil cosmid libraries) biosynthetic gene clusters containing AD and KS domain contigs that were abundant and/or highly expressed in tomato and cucumber root microbiomes.

**Figure 6.**
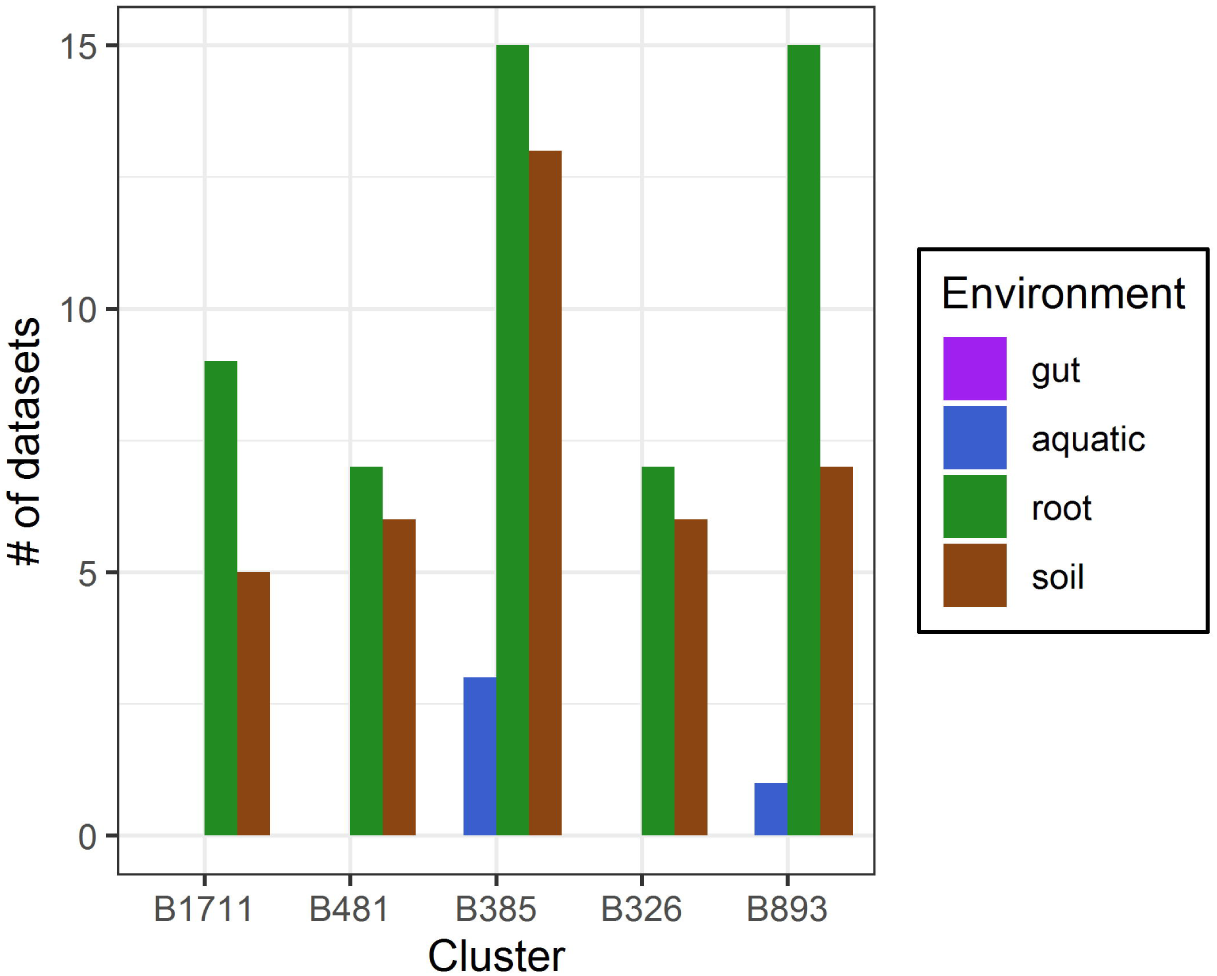
A) Comparison of B481 cluster and the related BGC in *S. cyaneogriseus*. Gray area indicate the aligned region. B) Structure prediction for PKS modules present on clone B893. Examples of known polyene antifungal agents are shown below.

**Table 2.** Predicted taxonomy and source dataset of recovered cosmids. UA-unassigned, only affiliated organisms with >80% identity are shown.

The five recovered BGCs were initially targeted due to their abundance in tomato and lettuce root datasets, suggesting a link to root ecosystems. To test this hypothesis, we assessed the abundance of the five BGCs in a large collection of publicly available shotgun metagenomes (20 metagenomes from each environment) from four distinct environments, targeting gut (animal and human), aquatic (freshwater and marine), soil (different soil types) and root-associated (various plant species) datasets. Our analysis demonstrated that the recovered BGCs are ubiquitous in most of the queried root samples and in some of the soil samples (**Fig. 7A**). Only B893 showed a significantly higher abundance in the root samples when compared with the soil samples (P<0.05, Wilcoxon test, **Fig. S5**). B893 was found in 16/20 of the root-associated datasets we examined (compared with 8/20 soil datasets, **Fig. 7B**). In contrast, none of the five BGCs were found in any of the gut microbiome communities analyzed, and very few were detected in the aquatic environments (**Fig. 7B**).

**Figure 7.**
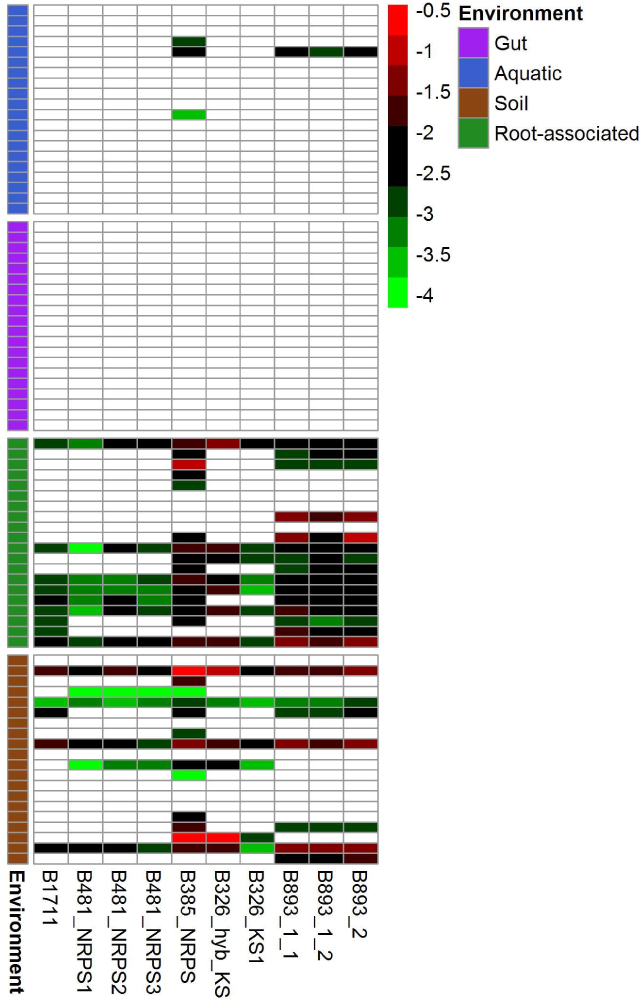
A) Relative abundance of clones-recovered AD and KS domains in shotgun metagenomes (n=20) from 4 environmental ecosystems-gut (animal and human), aquatic (freshwater and marine), soil and root-associated. Analysis was conducted using IMG BlastN feature with eValue<10-5 and identity >85%. Counts were normalized with rpoB and gyrB for each dataset. B) Presence/absence of each recovered gene cluster in the different environment. Only gene clusters where all AD/KS domains were found are considered present.

## Discussion

NRPs and PKs produced by root-associated microbial communities play an important role in plant root ecosystems (38, 39). Several studies have identified and characterized BGCs and/or associated metabolites in prominent plant growth promoting and biocontrol agents originally isolated from plant roots (40, 41). However, the large fraction of uncultivated bacteria in root ecosystems and the limitations associated with culturing bacteria encouraged us to examine the composition of genes encoding NRPS and PKS in root environments using culture-independent approaches. These methods have been applied before to understand SMs diversity and distribution in bulk soil (17, 26, 42, 43), but NRPSs and PKSs have not been comprehensively explored in root ecosystems. Furthermore, to the best of our knowledge, this is the first study to explore expression of secondary metabolites associated genes in root ecosystems.

Our results demonstrate a distinct composition of NRPSs and PKSs in plant (tomato and lettuce) roots relative to adjacent bulk soil; that these genes in root microbial communities are less diverse than those found in soil microbiomes; and that a fraction of these NRPSs and PKSs are highly expressed in root ecosystems. This is consistent with several previous studies showing that the phylogeny and functionality of root-associated microbial communities are significantly different from those in adjacent soil (44-46). It is well established that plants interact with soil bacterial communities through secretion of root exudates (47, 48), resulting in selection of specific microbial populations from the soil microbiome. This appears to be the case in the recruitment of plant-growth promoting rhizobacteria (PGPR), which are known to harbor specific SM encoding genes (49-51). Thus, while at this point we cannot infer the actual function of the highly abundant and expressed NRPs and PKs encoding genes in the root environment, we can infer that these NRPSs and PKSs likely play a role in various processes, *e*.*g*. competition and root colonization (52, 53).

While most of the detected SM-associated genes were too novel to link to any known metabolites, a few of the highly abundant, expressed and/or root-enriched NRPSs and PKSs were associated with known metabolites. Azalomycin F, for instance, found in both of the plant microbiome samples, is a polyketide with reported antifungal activity against a variety of phytopathogens, which is produced by different soil- and root-associated Streptomyces isolates (54, 55). Diaphorin, found only in the tomato root microbiome, is a pederin-analog known to be produced by the psyllid *Diaphorina citri* endosymbiont *Candidatus* Profftella armatura (Betaprotoebacteria), with potential antifungal and cytotoxic activity (56). In this regard, the described culture-independent approaches are a promising platform for identification of novel BGCs and for elucidating their roles in soil ecosystems and within the framework of drug discovery efforts, despite their current limitations (16, 17).

A large fraction of the highly abundant and highly expressed NRPSs and PKSs identified in this study were not assigned to, or had low identity (50-70%), when compared to previously characterized genes in the MIBiG repository. Recently, a previously unidentified hybrid NRPS/PKS gene cluster was found to be essential for Rhizoctonia suppression by an endophyte *Flavobacterium* sp. (57), highlighting the vast amount of root-associated SMs with unidentified functional roles, that undoubtedly play a pivotal role in bacterial-plant interactions.

We screened a large set of soil cosmid libraries with candidate sequences from our amplicon sequencing and metagenomic analyses that were enriched (relative to bulk soil), and/or highly abundant and expressed in tomato and lettuce roots, taking advantage of a unique culture-independent platform capable of extracting and analyzing long NRPSs and PKSs (27, 58). Five clones containing uncharacterized gene clusters with no known function, including two that were not associated with any known taxonomic group were identified. The fact that all of these BGCs were rather common in various root-associated environmental metagenomes, but rare or completely absent in other environments, suggests their potential importance in these habitats. While we can only speculate as to their synthesized metabolite’s actual activity, they were associated with bacterial groups well-known for their antimicrobial capacity. *Saccharohrix saharansis (*Actinobacteria), for instance, which contains a BGC closely related to clone B893, is a soil-dwelling bacteria known to produce an array of antimicrobials (59). This BGC likely encodes for a polyene substructure, often seen in antifungal agents, as it capable of directly disrupting the fungal membrane (60). Of particular interest is clone B481, associated with SM-associated genes from *Streptomyces cyaneogrisues*, known for its ability to produce the biopesticide Nemadectin (61). We speculate that this BGC may be associated with bacterial-fungal competition in the root ecosystems.

At the broader level, our results emphasize the need to look beyond basic descriptive diversity and composition information regarding SM capacity of microbial communities. The pipeline adopted in this study, where potentially important AD and KS sequences are first identified, followed by extraction of BGCs from cosmid libraries in order to identify potentially novel BGCs, has the advantages of being resource-efficient, while spawning deeper knowledge regarding potential important gene clusters. Future studies will focus on expressing these cosmid library BGCs in suitable hosts, enabling us to characterize their encoded metabolites and test their *in vitro* and *in planta* activity against various phytopathogens. Our results coincide with studies conducted in other plants, showcasing the yet unexplored diversity of NRPSs and PKSs (43, 62).

Overall, this study indicates that the root microbiome harbors a unique, diverse and potentially novel array of SM synthesizing genes, which are significantly different from those in the bulk soil microbiome. To enhance our current understanding, future research should focus on identifying additional factors shaping the occurrence and expression of SMs in the root microbiome (*e*.*g*., plant health, presence of phytopathogens and plant growth). This will undoubtedly help expose the ecological role of SMs in root ecosystems and provide a platform for drug discovery and novel and environmentally sustainable compounds for plant protection.

## Materials and Methods

**Fig. S6** present a conceptual description of the pipeline applied in this study.

### Amplicon sequencing, shotgun metagenomic and meta-transcriptomics analyses

Tomato soil and root samples and lettuce root samples were collected as previously described (63). Briefly, tomato (*Solanum lycopersicum*-Heinz 4107) and lettuce (*Lactuca sativa*-Romaine-Assaph) seedling were planted and grown for 42 days in a random block lysimeters experiment, at Lachish agricultural research station in Kiryat-Gat, Israel. Each sample type (soil, tomato roots and lettuce roots) was analyzed in triplicate samples from three different lysimeters (thus n=3 for each sample type). Each triplicate consisted of a composite sample collected from 2-4 plants or 3 soil samples, taken from the distant edges of the lysimeters and away from plant roots. As the same soil was used throughout the experiment, and given soil sample collection was taken for sufficient distance from growing plants, bulk soil from the tomato lysimeters served as a reference point for both tomato and lettuce soils. Previous study showed that they harbored almost identical microbial communities (Zolti et al. 2019). Soil samples were collected, frozen in liquid nitrogen on site, and stored at −80 °C for until further analysis. Root samples were collected intact, and soil particles were removed by shaking and briefly rinsing. The roots were then lightly dried and immediately frozen in liquid nitrogen on site and kept at −80 °C until processed. In this study, extracted DNA was used as template for NRPSs and PKSs PCR amplification using degenerated primers (A3F/A7R for AD-NRPS and degKS2F/degKS2R for KS-PKS domains, as previously reported (23). Resulted barcoded libraries were pooled and sequenced on an Iluumina MiSeq, employing V2 chemistry at the University of Illinois Chicago Sequencing Core (UICSQC). A total of 18 samples were sequenced, which included sampling location (tomato soil vs. tomato and lettuce roots) and SM family (AD and KS domains) with three replicates for each treatment. Resulting sequences were processed and demultiplexed using QIIME2 pipeline and the integrated DADA2 method (64). Exact Sequence Variants (ESVs) represented by fewer than 3 sequences were removed from downstream analyses. Raw amplicon sequences are available via the MG-RAST data depository, under mgm4862150.3. In addition, shotgun metagenome and metatranscriptome datasets of lettuce and tomato roots (n=3 for each dataset type, hence 6 for tomato and 6 for lettuce), previously generated and analyzed from same samples were also used for NRPSs and PKSs identification as described below (32). Shotgun sequences data is available via the NCBI Sequence Read Archive (SRA) data depository, with the project number PRJNA602301.

### Identification and annotation of NRPSs and PKSs

For chemical diversity analysis of NRPS and PKS gene clusters, the different datasets (ESVs generated via amplicon sequencing and metagenome/meta-transcriptome assembled genes) were aligned against the Minimum Information about a Biosynthetic Gene cluster repository (MIBiG, version August 6th, 2018). Only core NRPSs and PKSs genes (AD and KS domains) were included in the analysis. Alignment was performed using diamond blastx command line, with >50% amino acid sequence identity. To associate each NRPS or PKS hit with its potentially synthesized metabolite, e-Value <10^−40^ was used as a cut-off. Hits that did not pass this threshold were regarded as ‘unknown’.

For taxonomical annotation, sequences were aligned against the non-redundant (nr) Blast NCBI database, followed by lowest-common-ancestor (LCA) classification using MEGAN 6.15 Ultimate edition by taking the top 10% hits and filtering for a minimum score of 50 and maximum eValue of 0.01 (65). Conversion of gene identifiers to taxonomical path was done using the mapping files provided by MEGAN as of October 2016.

### Soil libraries amplicons generation screening

Metagenomic libraries were constructed as previously reported (66). Briefly, in each library, crude environmental DNA (eDNA) was extracted directly from field collected soil, gel purified, blunt ended, ligated into cosmid pWEB::TNC (Epicenter), packaged into lambda phage and transferred into Eco EC100 (Lucigen). Each library was expanded to contain 20 x106 unique cosmid clones with ∼30-45 kb eDNA (environmental DNA) inserts and then arrayed into 768 sub-pool (2×384-well plates) containing ∼25,000 unique cosmid clones per well. Each sub-pool was then stored as a glycerol stock for the clone recovery of interesting hits and as cosmid DNA to facilitate PCR-based screening. To generate an amplicon sequence database of NRPSs and PKSs, two set of degenerate primers (AD and KS) were applied to amplify the conserved region in Adenylation and Keto-synthase domains in the biosynthetic gene cluster: AD (forward) 5’-SATBTAYACSTCVGGHWCSAC-3’ and (reverse) 5’-CCANRTCNCCBGTSYKGTACA-3’; KS (forward) 5’-TGYTCSDSSTCGCTSGTSGCS-3’ (reverse) 5’-GTNCCSGTSCCRTGBGCYTCS-3’. The 5’ ends of the primers were augmented with Miseq sequencing adapters followed by unique 8 bp barcode sequences identifying the soil metagenome from which they were amplified. Amplicons were pooled as collections of 96 samples and cleaned using magnetic beads. Cleaned, pooled amplicons were used as template in a second PCR. Prior to sequencing, all PCR amplicons were quantified by gel electrophoresis and mixed in an equal molar ratio. The resulting pool was fluorometrically quantified with a HS D1000 ScreenTape and sequenced on an Illumina MiSeq instrument. Reads were debarcoded and trimmed to 240 bp. The reads from each sample were clustered using UCLUST (67) to generate the 95% tags.

### Recovery of BGC Clones from metagenomic library pools

The library well locations for target AD or KS domains were identified using well-specific barcodes incorporated in the degenerate primers (27). Specific primers with Tm value ∼60°C (18 to 20 bp) were designed to amplify each unique conserved sequence of interest. To recover the single cosmid clone from each library subpool, a serial dilution of whole-cell PCR strategy was used (17). Briefly, library glycerol stocks that contain target hits from eSNaPD analysis were inoculated into 3 ml LB broth (Kanamycin & Chloramphenicol) and grow overnight at 37 °C to confluence. The overnight cultured cells were diluted to 2000 CFU ml-1 calculated by OD600nm. 384 well plates were inoculated with 50 ul of the resulting dilution (600 CFU/well) with an Eppendorf epMotion 5075 liquid hander, grown overnight and screened using real-time PCR with touchdown PCR program, to identify wells containing target clones. Target positive wells were diluted to a concentration of ∼5 CFU ml-1 and the process repeated to identify new wells containing target clones. Five clone pools were then plated on solid agar plate and target single clones were identified by clone PCR.

### Analysis of Recovered Gene Clusters

Recovered single cosmid clones were miniprepped by QIAprep kit and sequenced using Miseq technology. M13-40FOR and T7 universal primers were utilized to sequence both ends of the insert sequences. Reads, amplicons and End sequences were assembled together to generate constant contigs using Newbler 2.6 (68). Fully assembled contigs were then analyzed using an in-house annotation script, consisting of open reading frame (ORF) prediction with MetaGeneMark, HMM Scan and BLAST search. The annotation script was developed using Python and is available in the open source repository: https://github.com/brady-lab-rockefeller/gene_annotation. Putative function and source organism of genes in the BGC were assigned based on the closest characterized gene in NCBI. KnownClusterBlast in antiSMASH 5.0 (69) was utilized to analyze the relationship between known characterized gene clusters and recovered BGC. The structure prediction of adenylation domain and keto-synthase domain in BGC was given by the antiSMASH prediction which employ three prediction algorithms, NPRSPredictor2, Stachelhaus code, and SVM prediction. These predicted building blocks were then utilized to predict final structure combined with known characterized BGC in cultured bacteria.

### Recovered clones search in environmental shotgun metagenomes

AD and KS domains from all five recovered gene clusters were searched against shotgun metagenomes from four different environments-animal and humans feces (gut), aquatic, soil and root-associated. We selected 20 Illumina-sequenced shotgun metagenomes from each of these ecosystems using the JGI IMG/MER advanced search option, followed by a blastn search using the IMG website online tool. Additional filtering was performed based on eValue threshold (10^−40^) and identity (>85%). For each hit, counts were normalized using gyrB and rpoB housekeeping genes counts (obtained via the IMG/MER platform). A relative abundance heatmap created using pheatmap R package (70). For gene clusters with more than one AD/KS domain, results are shown only for datasets that contained all cluster-belonging domains. Gene clusters presence/absence plot created using ggplot2 R package. Further information regarding selected metagenome datasets is shown in table S2.

### Statistical Analyses

Alpha (Simpson index) and beta (Bray-Curtis) diversity indices across environments (bulk soil vs. roots and soil vs. tomato vs. lettuce) were calculated using R package vegan (71). Variation of NRPS/PKS-associated genes was visualized by Principal Coordinate Analysis (PCoA) using same R package. To obtain this figure, we performed ordination on ESV count table (constructed by qiime2) using Bray-Curtis distances, followed by plotting using R ggplot2 package. Difference significance between groups was determined using vegan ANOSIM (Analysis of similarities) analysis.

Enrichment of ESVs between soil and tomato and lettuce roots and of NRPS/PKS-related sequences between roots shotgun metagenomes and metatranscriptomes was determined using DESeq2 (72). Only sequences with a corrected-adjusted P-value <0.1 (Wald test p-values corrected for multiple testing by Benjamini-Hochberg method) were chosen.

## Acknowledgments

We thank Avihai Zolti, Asaf Levi, Jonathan Friedman and Noa Sela for providing excellent technical and methodological help.

## Funding

This manuscript was supported by the US-Israel Binational Agricultural Research and Development Fund (BARD) Grant #: IS-5177-19F.

## Supplemental material legend

**Figure S1.** Alpha diversity (1-Simpson similarity index) of soil and root (tomato and lettuce) samples. **A and B**-AD sequences (NRPS). **C and D**-KS sequences (PKS). Asterisks represent statistical significance between samples (by Tukey HSD, *- pValue<0.001, **- pValue<0.0001, NS- not significant).

**Figure S2**. Relative abundance of NRPS (A) and PKS (B) associated metabolites based on MIBiG annotations. Only AD and KS amplicon hits to reference MIBiG sequences over >50% amino acid identity and e-Value<10^−40^ were included. Counts were normalized and log10-transformed. Only associated metabolites that were present in >2 different samples are shown.

**Figure S3**. The fifty most abundant NRPSs and PKSs in tomato (A) and lettuce (B) root microbiomes extrapolated from shotgun metagenomic data. Relative abundance represent the mean of three repeats per sample. CV (coefficient of variance) was determined as STD/Mean per sequence. Green-circled points represent sequences that are highly abundant in the corresponding meta-transcriptome datasets per sample type indicating that these are also highly expressed.

**igure S4**. Mined NRPSs and PKSs from amplified AD and KS domains in tomato and lettuce roots, tomato metagenome and lettuce metagenome. AS-amplicon sequencing, MG-metagenome. Numbers in circles represent number of unique MIBiG-derived genes found within each dataset, while numbers in parentheses represent percent of unique genes from total.

**Figure S5. Relative abundance of recovered gene clusters in root-associated and soil metagenomic samples**. Abundance was normalized to gyrB and rpoB housekeeping genes within each dataset (n=20 per environment). Statistical difference was calculated using Wilcoxon non-parametric test. *= p<0.05, NS-not-significant

**Figure S6**. Study conceptual pipeline scheme

**Table S1**. Amplicon sequencing processing information.

**Table S2**. Public shotgun metagenome of microbial communities from four different ecosystems (animal and human gut, aquatic, soil and root-associated).

## Notes

### Competing Interest Statement

The authors have declared no competing interest.

### Summary of Updates

Figures content and order were changed, MS content was revised, supp. information was changed.

