## Supplementary figures and images for "Elucidating the diversity and potential function of nonribosomal peptide and polyketide biosynthetic gene clusters in the root microbiome"

### Supplemental Figure 1

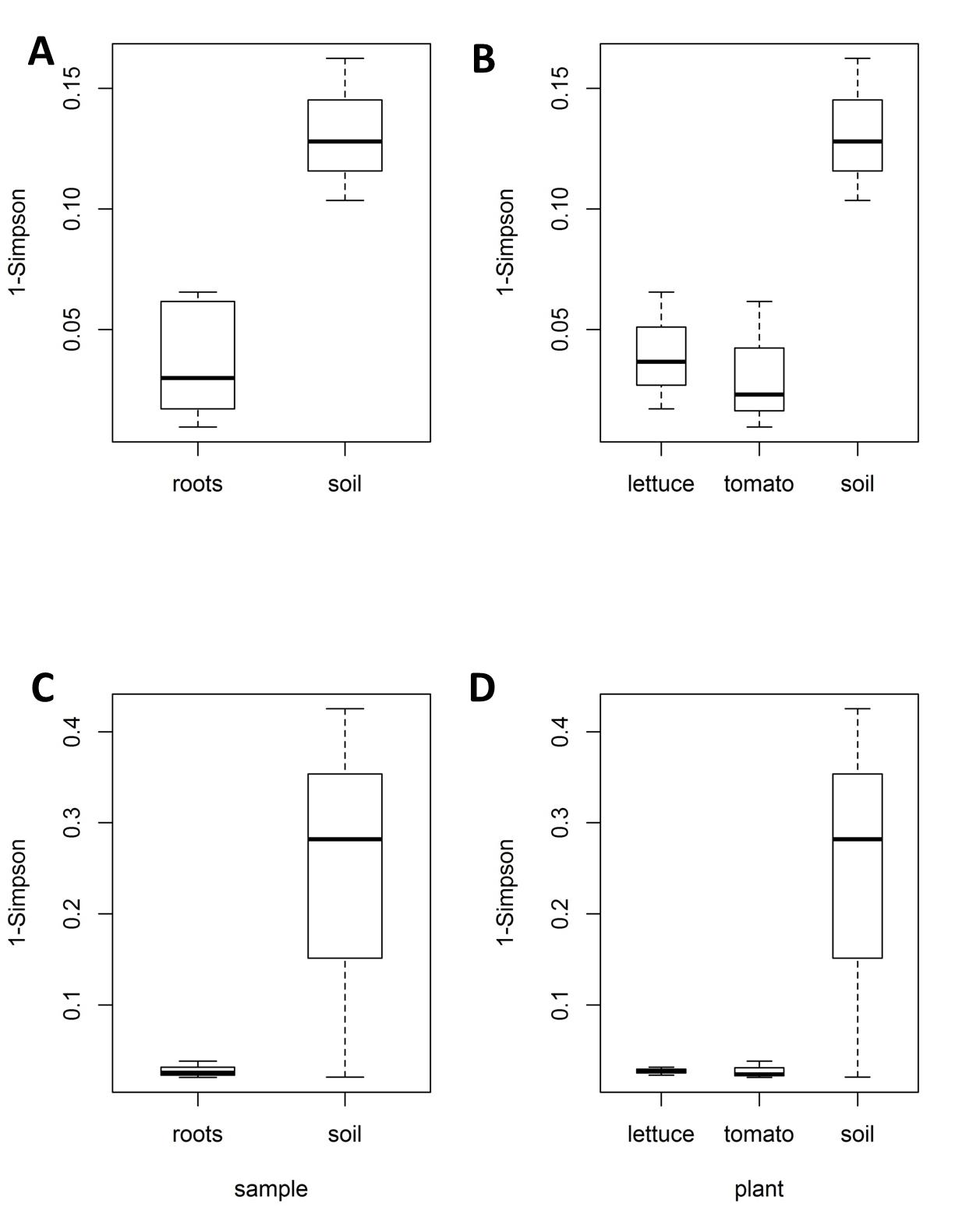

### Supplemental Figure 2

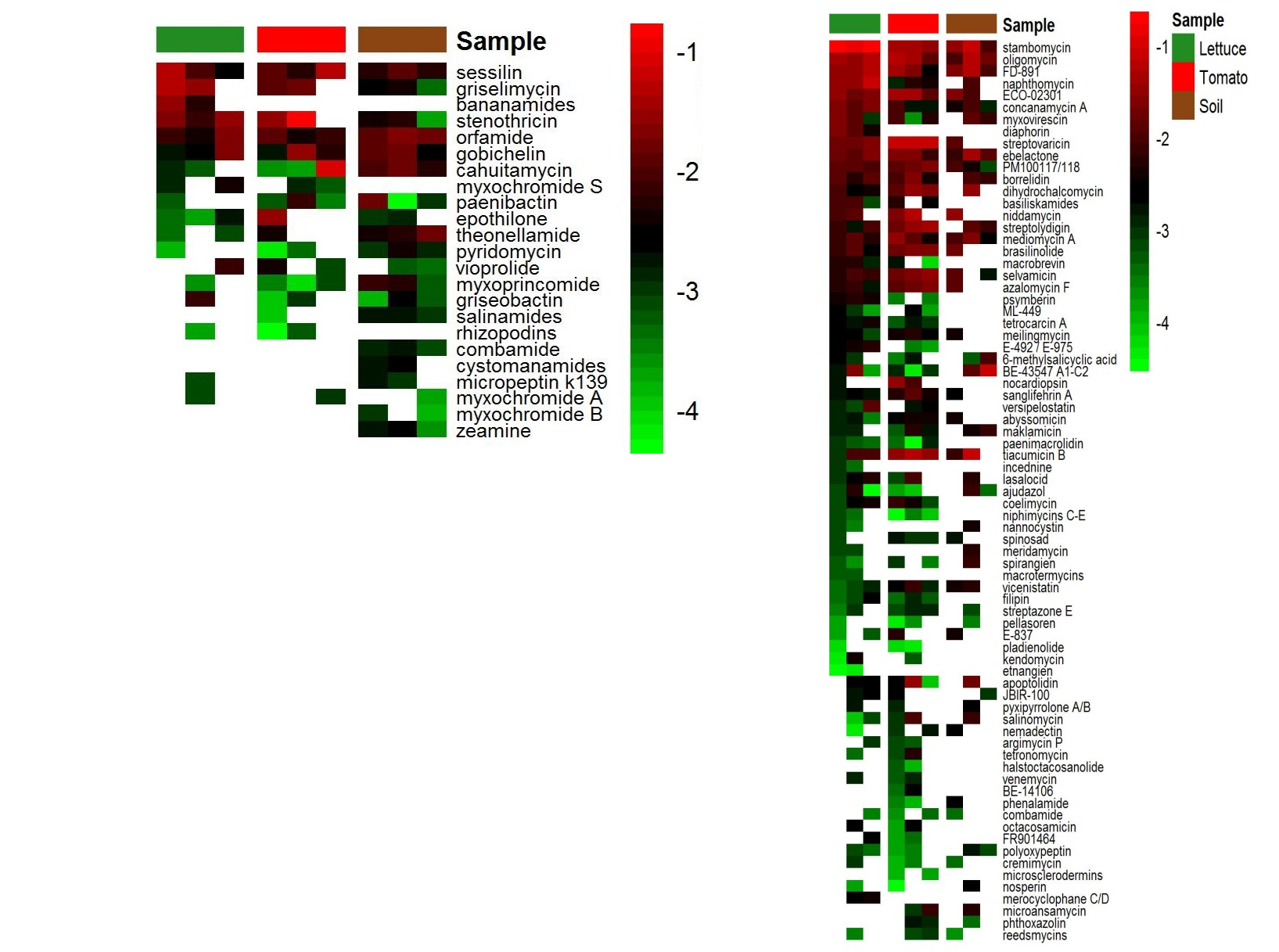

### Supplemental Figure 3

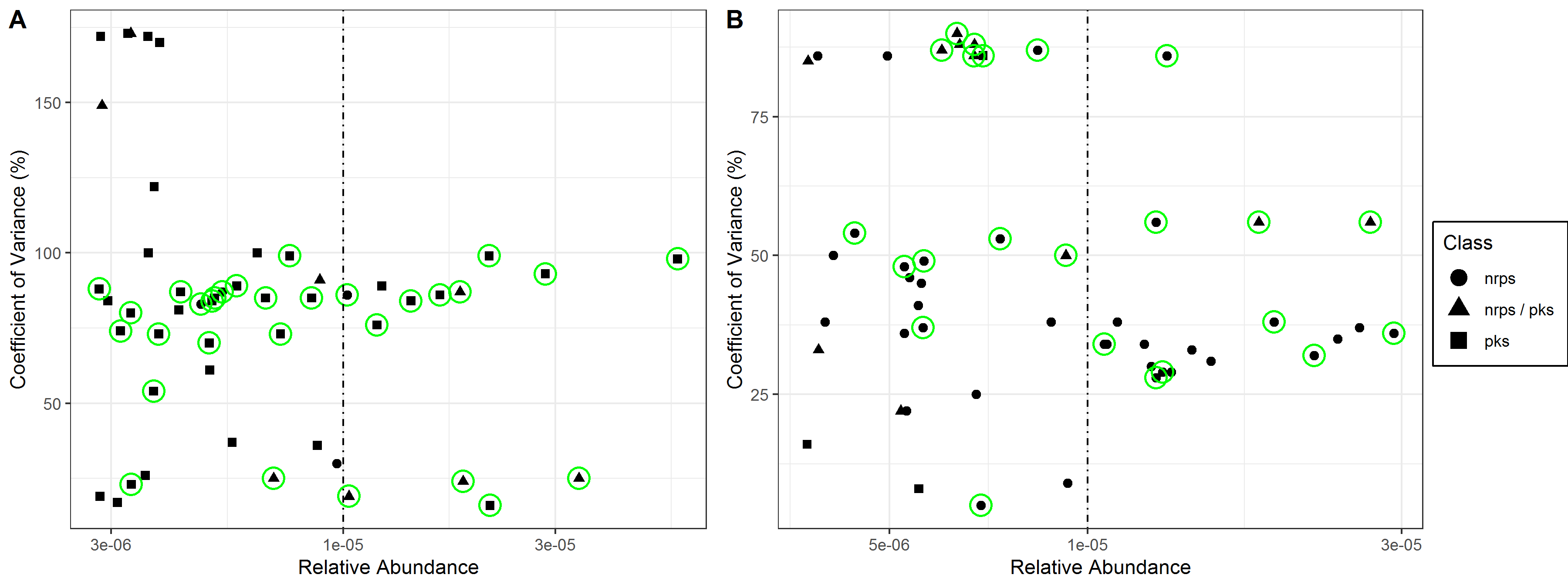

### Supplemental Figure 4

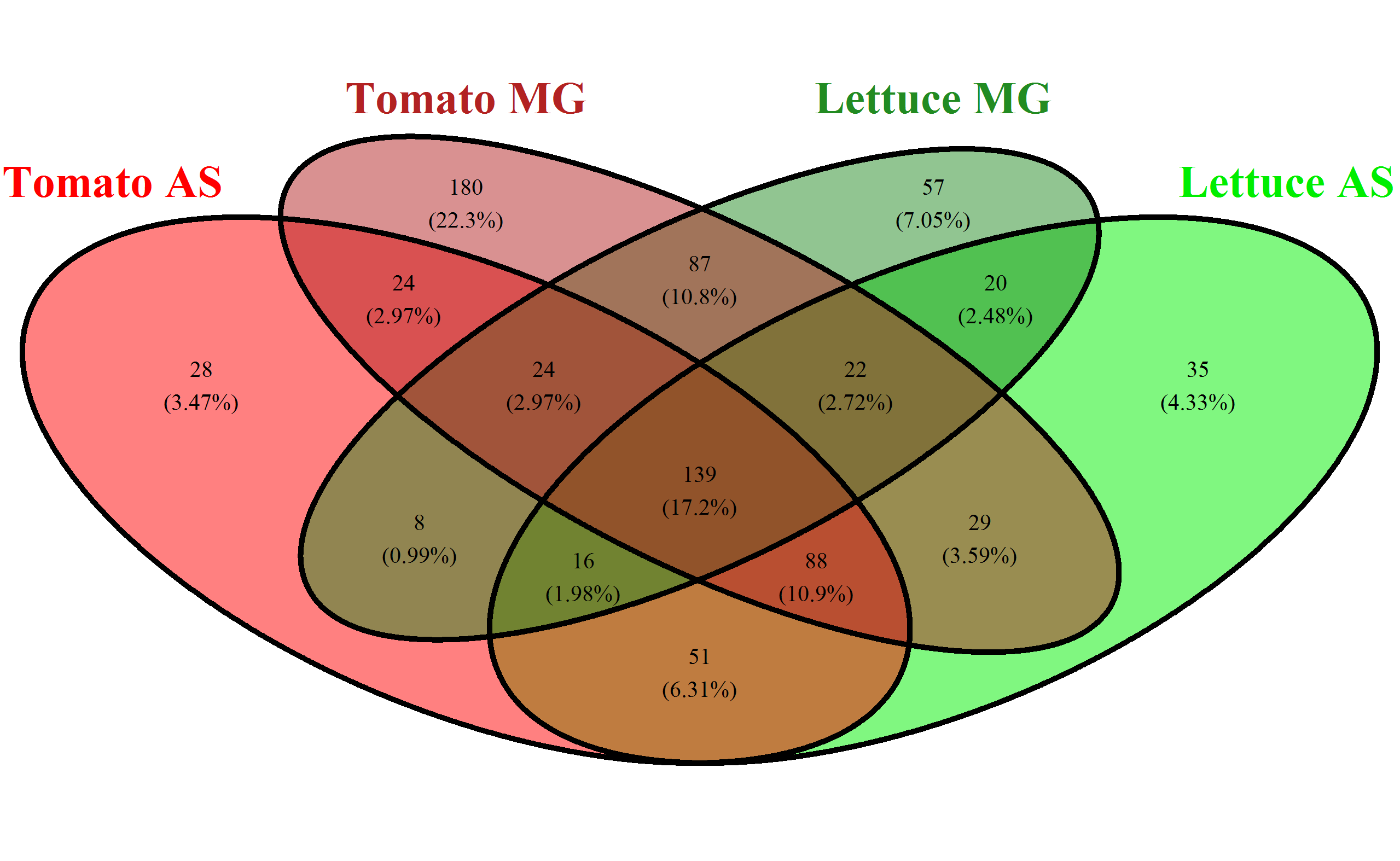

### Supplemental Figure 5

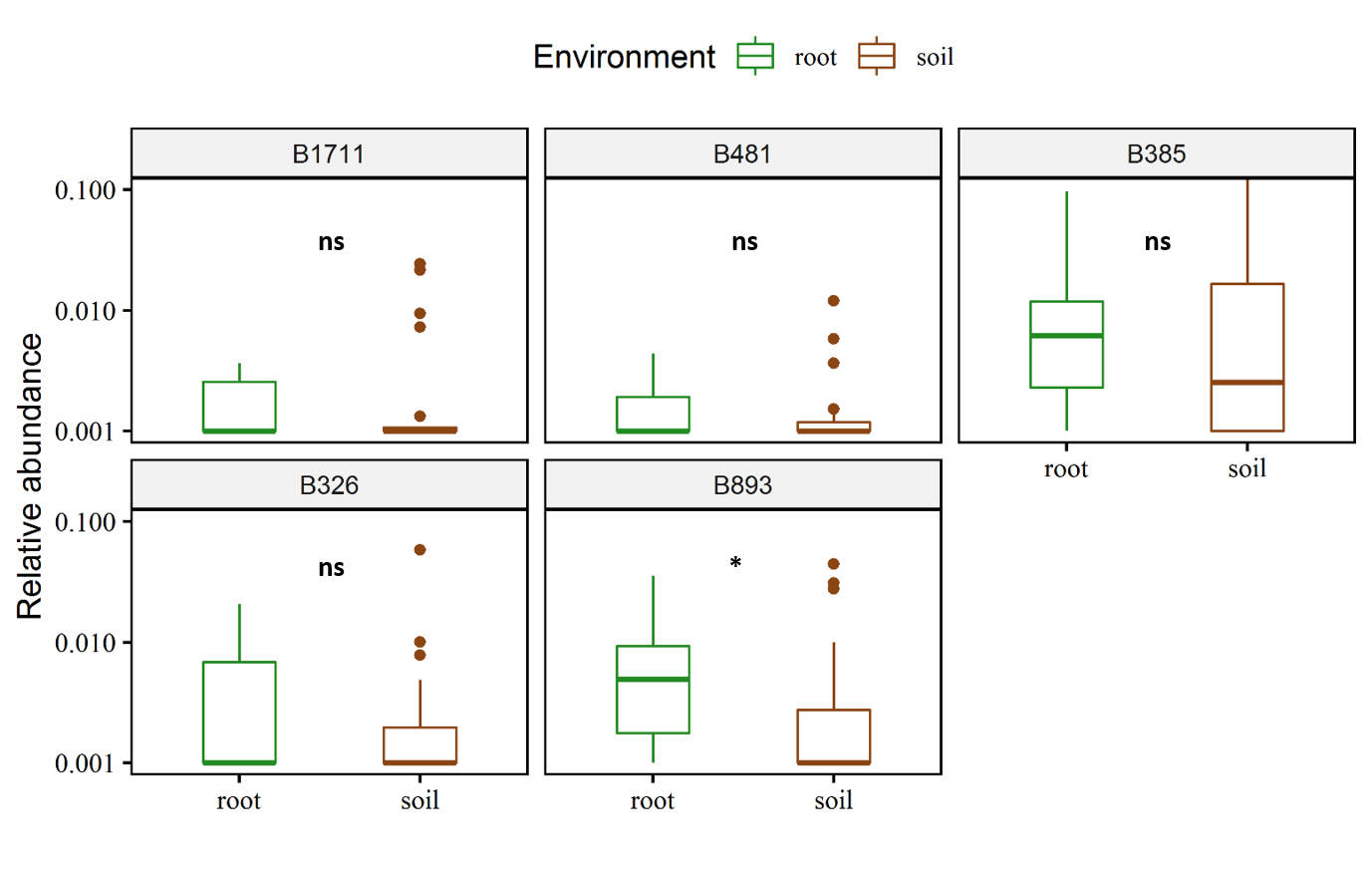

### Supplemental Figure 6

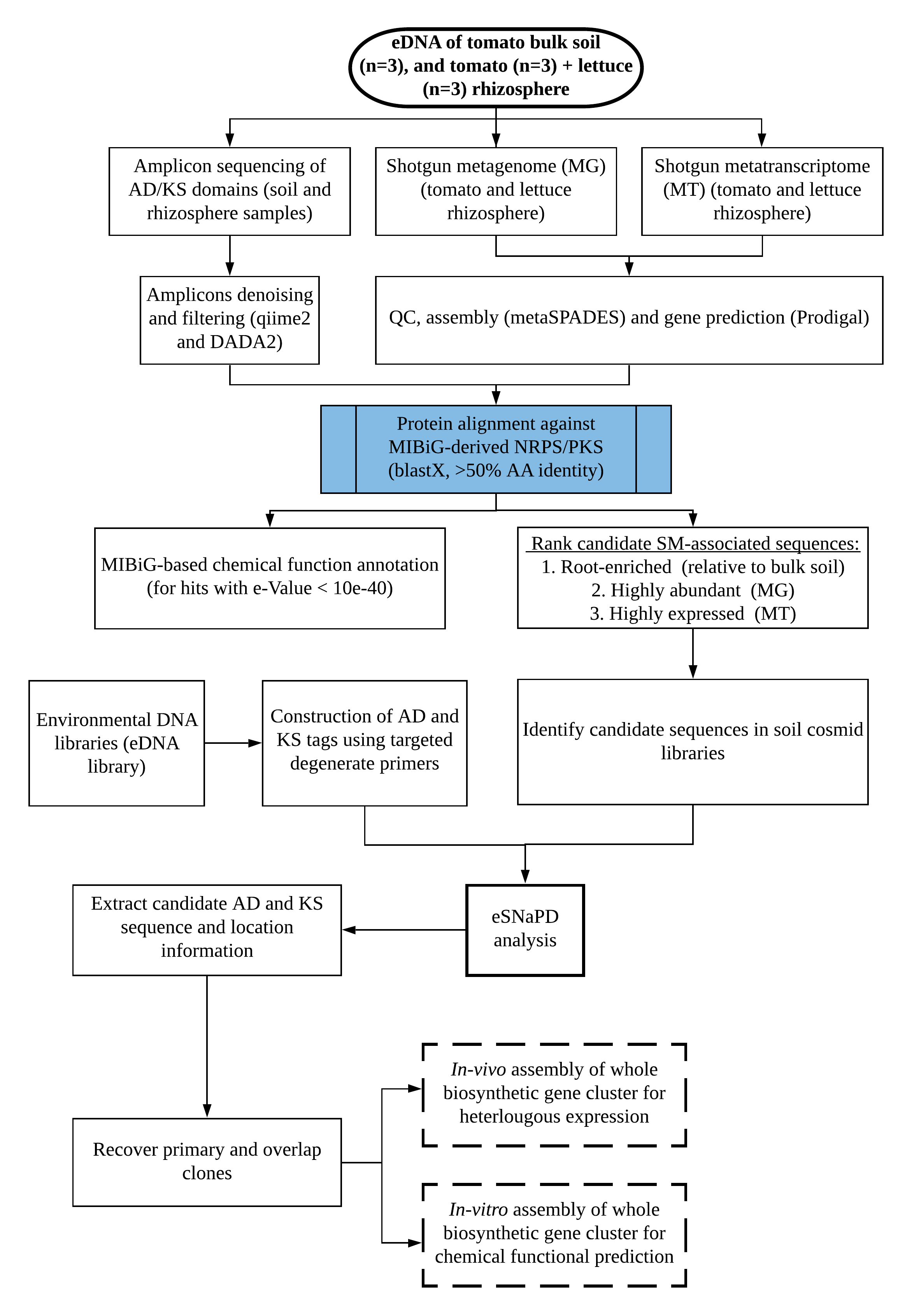
